# Sequential increase of PHGDH expression with Alzheimer’s pathology and symptoms

**DOI:** 10.1101/2022.02.15.480384

**Authors:** Xu Chen, Riccardo Calandrelli, John Girardini, Zhangming Yan, Zhiqun Tan, Xiangmin Xu, Annie Hiniker, Sheng Zhong

## Abstract

We report consistent increases in phosphoglycerate dehydrogenase (PHGDH) expression in mouse models of Alzheimer’s disease (AD) (3xTg-AD) and related tauopathy (PS19), particularly in hippocampal astrocytes. Human single-cell RNA-sequencing data reveal a sequential increase of PHGDH expression in people with no, early, and late AD pathology, which is corroborated by protein mass spectrometry and immunohistochemical analyses of three independent cohorts. A sequential increase of PHGDH expression also correlates with increasing clinical AD symptoms and worsening cognitive decline in patients. The consistent increase of PHGDH expression in six AD cohorts (Mayo, ROSMAP, Mount Sinai, Baltimore, Amsterdam, and UCSD/UCI) corroborates with the recent report of PHGDH extracellular RNA in blood plasma as a candidate biomarker for early diagnosis of AD and offers a caution to the suggested use of L-serine as a potential therapy of AD.

## Introduction

Phosphoglycerate dehydrogenase (PHGDH) is required for the synthesis of serine, a modulator of synaptic plasticity (Guercio and Panizzutti, 2018). Human mutations in PHGDH can cause abnormal brain development due to serine deficiency while conditional knockout of PHGDH in the mouse hippocampus impairs synaptic plasticity and spatial memory, in part by diminishing NMDA receptor (NMDAR)-evoked neuronal immediate early genes (Le Douce et al., 2020; Neame et al., 2019; Yang et al., 2010). Given the importance of synaptic pathophysiology to Alzheimer’s disease (AD), these results raise the possibility that abnormalities in PHGDH expression could contribute to AD pathogenesis. A missing key to this question is the direction of change of either serine level or PHGDH expression levels in AD. Despite extensive efforts, changes in L- or D-serine’s level have not been unequivocally correlated with AD, making it particularly important to assess changes in PHGDH expression that may occur in AD.

We read with great interest the recent publication in *Cell Metabolism* reporting that “expression of PHGDH is reduced in the AD brain” (Le Douce et al., 2020). Leveraging this result to bridge the gap between serine/PHGDH deficiency and AD pathogenesis, the authors suggested oral L-serine as “a ready-to-use therapy for AD” (Le Douce et al., 2020). In contrast to this report, our previous meta-analysis of the National Institute on Aging’s Accelerating Medicines Partnership-Alzheimer’s Disease (AMP-AD) consortium’s RNA-seq datasets revealed a consistent increase of PHGDH mRNA expression in AD in five brain regions from Mayo, ROSMAP, and Mount Sinai cohorts (Yan et al., 2020). Consistent with our previous findings, and in contrast to the work of Le Douce et al., here we report that PHGDH mRNA and protein levels are increased in the brains of two mouse models of AD and/or tauopathy, and are also progressively increased in human brains with no, early, and late AD pathology, as well as in people with no, asymptomatic, and symptomatic AD.

## Results

### PHGDH expression is increased in 3xTg-AD mice

We report an increase in PHGDH expression in 3xTg-AD mice, an AD model that develops both amyloid-β and tau pathology. First, we reanalyzed an RNA expression dataset (GSE144459) of 3xTg-AD mice. Hippocampal PHGDH mRNA level in 3xTg-AD mice is not different (Bonferroni-corrected p-value = 0.682, T test) from non-transgenic (Non-Tg) controls at age 3 months, when AD pathology has not developed, but is higher (Bonferroni-corrected p-value = 0.015, T test) than Non-Tg controls at 12 months.

Next, we repeated Le Douce et al.’s immunofluorescent confocal analysis. To ensure reproducibility, we obtained hippocampal sections from five pairs of 6-month-old female 3xTg-AD and Non-Tg controls. Hippocampal PHGDH immunostaining was increased in 3xTg-AD compared to Non-Tg (p-value = 0.0039, n=5/group, permutation test) (Figure S1A, B). Our five pairs of mice were raised by four different labs, strongly arguing against lab-to-lab variation. Additionally, we observed an increase in GFAP expression in 3xTg-AD compared to Non-Tg (Figure S1A). Thus, hippocampal PHGDH expression is increased in 3xTg-AD mice compared to non-transgenic controls.

### PHGDH expression is also increased in a human tau transgenic mouse model PS19

Recognizing that no single mouse model can fully recapitulate human AD, we asked if our observation in 3xTg-AD mice could be reproduced in PS19 mice, a model for neurofibrillary tangle formation in AD that is independent of amyloid-β. We assessed PHGDH protein level changes via immunohistochemistry and western blots. In PS19 mice, hippocampal PHGDH exhibited significant colocalization with GFAP, confirming astrocyte expression of PHGDH (Figure S1C) (Le Douce et al., 2020; Yang et al., 2010). Compared to non-TG littermate controls, PHGDH protein levels were increased in the hippocampus of 10-month-old PS19 mice (p-value for western blots < 0.0001, unpaired T test, n=8 for PS19 and n=7 for Non-Tg) (Figure S1C-E), suggesting that expression of a human mutant tau transgene is sufficient to induce hippocampal PHGDH expression.

### Single-cell transcriptomes reveal a sequential increase of PHGDH with AD pathology

We report a sequential increase of PHGDH mRNA expression with AD pathology in humans. We reanalyzed 80,660 single-nucleus transcriptomes from 24, 15 and 9 individuals with no, early, and late AD pathology (Mathys et al., 2019). Consistent with reported astrocyte expression of PHGDH (Yang et al., 2010), PHGDH was detected in approximately 10-25% of astrocytes (Ast), oligodendrocytes (Oli), and oligodendrocyte precursor cells (Opc), and less than 4% of excitatory (Ex) and inhibitory (In) neurons, endothelial cells (Ec), and microglia (Mic) (Figure S1F). PHGDH exhibits a sequential increase of expression from no to early and to late pathology in astrocytes (p-value = 0.0016, ANOVA controlling for sex. No multiple hypothesis testing is involved.) (Figure S1F). The fraction of PHGDH-expressing astrocytes does not increase from no to early pathology (green vs. blue Ast columns, Figure S1G), suggesting that the early pathology-associated PHGDH expression increase (green vs. blue Ast columns, Figure S1F) is due to an increase in the expression level per cell rather than an increase in the proportion of PHGDH-expressing cells.

### Sequential increases of PHGDH protein with AD pathology and symptoms

We identified sequential increases in PHGDH protein expression with AD pathology and symptoms. We reanalyzed two human mass spectrometry datasets (Hondius et al., 2016; Seyfried et al., 2017). PHGDH protein levels increase with increasing Braak stage in an Amsterdam cohort of 40 individuals (p-value = 0.013, ANOVA controlling for sex, age, and postmortem delay (PMD)) (Figure S1H, I). PHGDH expression also increases from controls to asymptomatic AD (ASYMAD, these people exhibited no clinical symptom, but have autopsy identified pathology) and to AD (people with symptoms and autopsy confirmed pathology) in dorsolateral prefrontal cortex from a Baltimore cohort of 42 individuals (p-value = 0.040, ANOVA controlling for sex, age, and PMD) (Figure S1J, K). These data suggest an increase in PHGDH protein expression during both pathological and symptomatic progression of AD.

To validate the mass spectrometry results, we carried out PHGDH immunostaining on 21 hippocampal samples from age-matched cases and controls. We compared 10 samples at Braak stages 0-3 (negative-early group (NE)) with 11 samples at Braak stage 4-6 (advanced AD group (AD)). PHGDH immunostaining is higher in AD hippocampi than in NE hippocampi (p-value < 0.02, ANOVA controlling for sex) (Figure S1L, M), confirming the mass spectrometry result from the Amsterdam cohort. Next, we separately analyzed CA1, CA3, and dentate gyrus (DG). PHGDH immunostaining is increased in all three regions in AD compared to NE. This AD-associated increase is pronounced in CA1 and CA3 (p-value < 0.006, ANOVA controlling for sex) (Figure S1N) but not significant in DG (p-value = 0.72, ANOVA controlling for sex). These data corroborate with the AD-pathology-associated increase of PHGDH expression.

To validate the correlation of PHGDH protein level with AD’s symptomatic development, we utilized the Dementia Rating Scale-2 (DRS), a metric of a patient’s overall level of cognitive functioning (larger DRS indicates better overall cognitive ability). Ten of our analyzed hippocampal samples have corresponding DRS. In these samples, PHGDH immunostaining decreases as the donor’s DRS increases (p-value < 0.0001, ANOVA controlling for sex). Next, we separately analyzed CA1, CA3, and DG. PHGDH immunostaining decreases in each region as the DRS increases (p-value < 0.0015 in CA1 and CA3 (Figure S1O), p-value < 0.02 in DG). These data suggest an increase of hippocampal PHGDH expression as a patient’s overall cognitive function declines, corroborating the AD-symptom-associated increase of PHGDH in the Baltimore cohort.

## Discussion

The reproducible increase of PHGHD expression during both pathological and symptomatic developments of AD suggests different underlying mechanisms between PHGDH deficiency and AD. Thus, we feel “oral L-serine as a ready-to-use therapy to AD” (Le Douce et al., 2020) warrants precaution. This is because despite being a cognitive enhancer, some evidence suggests that long-term use of “D-serine contributes to neuronal death in AD through excitotoxicity” (Guercio and Panizzutti, 2018). Furthermore, D-serine, as a coagonist of NMDAR, would be expected to oppose NMDAR antagonists, which have proven clinical benefits in treating AD.

The opposite results between our and Le Douce et al.’s analyses may be attributable to the difference in the PMD of the human samples. Le Douce et al.’s AD samples had longer PMD than their control samples because five of their six controls (83%) as compared to only five out of their 15 AD samples (33%) had less than 30 hours of PMD (Le Douce et al.’s Table 1). Considering that human PHGDH protein is sensitive to protease cleavage at room temperature, PMD-related protein degradation may explain the lower PHGDH levels in Le Douce’s AD samples. The longest PMD in our reanalyzed mass spectrometry datasets is 30 hours (Hondius et al., 2016; Seyfried et al., 2017), and the PMD distributions in AD and control are not statistically different (Figure S1I,K). The longest PMD of the human hippocampal samples used in our immunostaining analysis is 18 hours.

Our results lead to a hypothetic model of positive feedback between increased astrocyte PHGDH expression and excitotoxicity. In this model, an increase in PHGDH expression in brain astrocytes leads to an increase in the basal level of NMDAR-dependent synaptic activities (Neame et al., 2019), with an increase in the probability of initiating an amyloid-β-dependent “vicious cycle of neuronal hyperactivation”, eventually leading to excitotoxicity (Zott et al., 2019). Excitotoxicity and pathological protein deposition induce astrogliosis, which further increases astrocyte PHGDH level and thus creates an ongoing vicious cycle of increased astrocyte PHGDH, excitotoxicity, and astrogliosis. If this model is correct, suppression of astrocytic PHGDH expression may relieve excitotoxicity. Consistent with this idea, reduction of D-serine by knocking out serine racemase, the L-serine to D-serine conversion enzyme, has a protective effect on amyloid-β toxicity in a mouse model (Inoue et al., 2008).

Our new results also support the prior observation of an AD-associated increase in PHGDH extracellular RNA (exRNA) in human plasma (Yan et al., 2020). Further, they help explain why the longitudinal increase of PHGDH exRNA can predict AD’s clinical diagnosis (Yan et al., 2020). Together, these data nominate circulating PHGDH exRNA as a possible diagnostic biomarker of late-onset AD.

## Acknowledgment

This work is partially funded by NIH grant UG3CA256960 and Kruger Research Award. The authors thank the University of California Alzheimer’s Disease Research Centers (UCI ADRC, UCSD ADRC) for providing the postmortem tissue samples. The authors thank Drs. Elizabeth Head, James Brewer, Robert Rissman, Edward Koo, Douglas Galasko, and Timothy Gahagan for samples or useful discussions. UCI ADRC is funded by NIH Grant P50AG16573. UCSD ADRC is funded by NIH grant P30AG062429.

## SUPPLEMENTAL FIGURES

**Figure S1.**
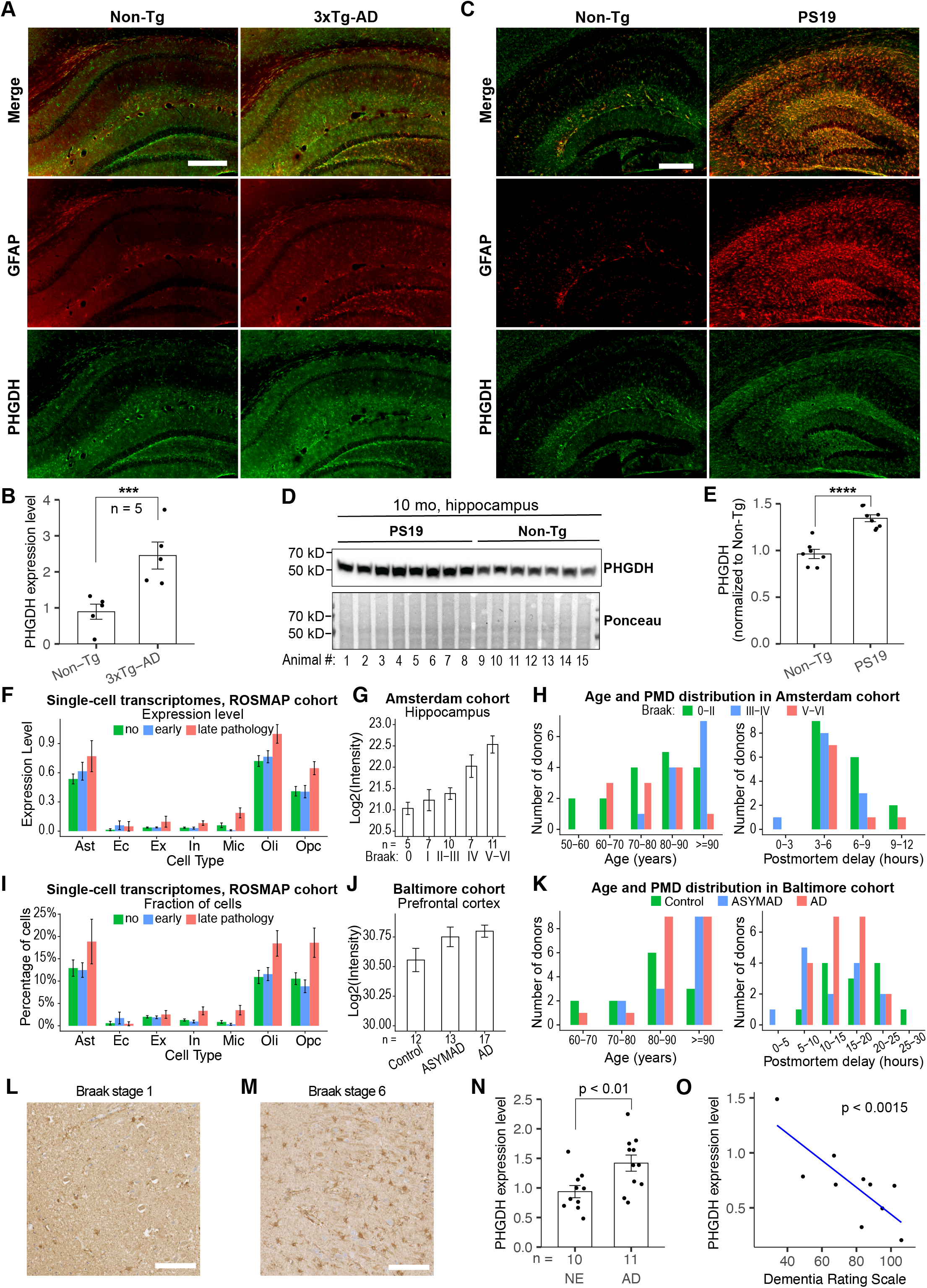
Comparison of PHGDH expression in AD and control brains. (**A**) Immunostaining of 6-month non-transgenic control (Non-Tg, left) and 3xTg-AD (right) hippocampus with PHGDH (green) and GFAP (red) antibodies. Scale bar: 300 μm. (**B**) Quantification of hippocampal PHGDH immunostaining (y axis) in 3xTg-AD and Non-Tg mice (columns). n=5 in each group. (**C**) Immunostaining of 10-month non-transgenic littermate control (Non-Tg, left) and PS19 (right) hippocampus with PHGDH (green) and GFAP (red) antibodies. Scale bar: 300 μm. (**D**) Immunoblot of PHGDH and Ponceau staining in the hippocampal lysates of 10-month PS19 and Non-Tg mice. (**E**) Quantification of PHGDH western blots as in (D), n=7-8/group. (**F-G**) PHGDH expression level (F) and the fraction of PHGDH-expressing cells (G) in people with no (green), early (blue), and late (pink) AD pathology in each cell type (column). Expression level was calculated by the default method in Seurat, i.e., log(10000×UMI/cell_total_UMI+1). The increase of expression in early pathology in Ast (Ast column, F) does not involve in an increase of fraction of PHGDH-expressing cells (Ast column, G). (**H**) A sequential increase of hippocampal PHGDH protein levels (y axis) with respect to Braak stages (x axis) in the Amsterdam cohort. (**I**) Age and PMD distributions of the Amsterdam cohort. (**J**) A stepwise increase of PHGDH protein levels in people with no symptom nor pathology (Control), pathology but no symptom (ASYMAD), and both pathology and symptoms (AD) in the Baltimore cohort. (**K**) Age and PMD distributions of the Baltimore cohort. (L-O) Hippocampal PHGDH immunostaining analysis. (L-M) Representative images from a Braak stage 1 (**L**) and a Braak stage 6 sample (**M**). Scale bar = 100 μm. (**N**) Quantification of PHGDH immunostaining. Average PHGDH expression in CA1 and CA3 (y axis) in every advanced AD (AD column, Braak stages 5-6, n=11) and negative-early (NE) sample (NE column, Braak stages 0-3, n=10) (columns). p: the p-value from ANOVA analysis. (**O**) Scatter plot of every sample’s average PHGDH expression level in CA1 and CA3 (y axis) vs. the donor’s Dementia Rating Scale (x axis). n=10. p: the p-value from ANOVA analysis.

## SUPPLEMENTAL METHODS

### Immunohistochemistry analysis of mouse brain

Anesthetized mice were transcardially perfused with 0.9% saline. Mouse brains were removed and fixed in 4% paraformaldehyde for 48 h. Fixed brains were cryopreserved in 30% sucrose in PBS for at least 2 days, and coronal brain sections (30 μm in thickness) were obtained with a sliding microtome. For immunostaining, floating brain sections were permeabilized and incubated in blocking solution (10% normal goat serum in 0.3% Triton X-100 PBST) at room temperature for 1 h. Sections were then incubated with primary antibodies, including rabbit anti-PHGDH (Proteintech 14719-1-AP, 1:500) and mouse anti-GFAP (CST #3670, 1:300). After overnight incubation, the sections were incubated with secondary antibodies, including fluorescein-labeled goat anti-mouse IgG (1:500, Vector Laboratories 1:1000, Life Technologies). Images were acquired by Keyence BZ-X710 microscope and analyzed with ImageJ software.

### Homogenization of brain tissue and immunoblotting

Mouse brain tissues were homogenized in RIPA buffer containing protease inhibitor cocktail (Sigma), 1 mM phenylmethyl sulfonyl fluoride, phosphatase inhibitor cocktail (Sigma), 5 mM nicotinamide (Sigma) and 1 μM trichostatic-A (Sigma), and sonicated after homogenization. Lysates were centrifuged at 14,000 RPM at 4 °C for 15 min. Supernatants were collected and protein concentrations were determined by the Pierce assay (Thermo Fisher). The same amount of protein was resolved on a 4-12 % SDS-PAGE gel (Invitrogen), transferred to PVDF membrane (Bio-Rad), and probed with appropriate antibodies. Bands in immunoblots were visualized by enhanced chemiluminescence (Pierce) and quantified by densitometry and ImageJ software (NIH). Representative blots from same gel/membrane are shown and compared in the same figure.

### Immunohistochemistry analysis of human brain samples

Paraffin-embedded brain slices pre-mounted on slides were heated in a lab oven for 30 min at 50C and de-paraffinized in xylene baths for 10 min each, followed by rehydration by sequential incubation with 100%, 90%, 70% EtOH, 5 min for each bath. Slides underwent antigen retrieval with Citrate buffer pH6 solution in a pressure cooker on low pressure for 10 min. The slides were blocked in 2.5% normal horse serum for 1 h at room temperature and the primary rabbit polyclonal PHGDH antibody was diluted at 1:500 or 1:2000 (in different cohorts of samples) in diluted normal horse serum and applied to the slides overnight in a humidified chamber at 4C. Signal amplification and detection were performed using Vector Laboratories VECTASTAIN® Elite ABC-HRP Kit, Peroxidase, R.T.U. (Universal) (PK-7200) and ImmPACT® DAB Substrate, Peroxidase (HRP) (SK-4105) according to the manufacturer’s instructions. Then slides were counterstained with hematoxylin for 3 min, washed, dehydrated, and coverslipped. Slides were scanned using a NanoZoomer Slide Scanner imager.

### Analysis of single-nucleus RNA-seq data

Single-nucleus RNA-seq (snRNA-seq) data were downloaded from the Synapse Portal (Synapse ID: syn2580853) and analyzed with the R package Seurat (v3). The nuclei with either more than 8,000 or less than 200 detected genes or with more than 10% of reads mapped to the mitochondrial genome were filtered out. The following marker genes were used: NRGN for excitatory neurons (Ex), GAD1 for inhibitory neurons (In), AQP4 for astrocytes (Ast), MBP for oligodendrocytes (Oli), CSF1R and CD74 for microglia (Mic), VCAN for oligodendrocyte progenitor cells (Opc), FLT1 for endothelial cells (Ec), and AMBP for pericytes (Per). Log-transformed expression levels were used for making a binary call (low- or high-expression) for every marker gene in every nucleus. A Gaussian Mixture Model (GMM) with two components was applied to the expression level of each marker gene (or the average expression of two marker genes for microglia) across the nuclei to assign each nucleus to one of the two components, namely the high- and the low-expression components.

Control and AD samples were first separately normalized with “sctransform” and then integrated into one dataset based on Seurat-selected anchor genes. This integrated dataset was used for clustering analysis. Clusters of nuclei were identified by Seurat based on shared nearest neighbor (SNN) using the first 10 principal components as input (resolution = 0.5). Each cluster of nuclei was assigned to a cell type based on the marker gene exhibiting the highest level of enrichment (Fisher’s exact test) in that cluster.

### Quantification of immunostaining images

Immunostaining images were analyzed using ImageJ with the following workflow. (1) The image format was changed to 16 bit. (2) Outlier pixels were removed. (3) Particle analysis was done with the threshold of size above 120 pixel square and circularity between 0.2 and 1. (4) PHGDH stained percent area was calculated as the ratio of the number of pixels within particle analysis detected spots and the total number of pixels (pixels under spots / total pixels). Standard neuroanatomically defined regions were assessed microscopically by board-certified neuropathologists. The ANOVA model for comparison of advanced AD (AD) vs. negative-early (NE) is: PHGDH percent area ~ disease + region + sex + site, where disease is AD or NE, region is CA1, CA3, or dentate, site is UCSD or UCI. The ANOVA model for DRS analysis is: DRS ~ PHGDH percent area + sex + region, where region is CA1, CA3, or dentate. There is no “site” variable in the ANOVA model for DRS analysis because all the samples with available DRS information were from UCSD.

